# MetaFetcheR: An R package for complete mapping of small compound data

**DOI:** 10.1101/2021.02.28.433248

**Authors:** Sara A. Yones, Rajmund Csombordi, Jan Komorowski, Klev Diamanti

## Abstract

**Motivation:** Small-compound databases contain large amount of information for metabolites and metabolic pathways. However, the plethora of such databases and the redundancy of their information lead to major issues with analysis and standardization. Lack of preventive establishment of means of data access at the infant stages of a project might lead to mislabelled compounds, reduced statistical power and large delays in delivery of results.

**Results:** We developed MetaFetcheR, an open-source R package that links metabolite data from several small-compound databases, resolves inconsistencies and covers a variety of use-cases of data fetching. We showed that the performance of MetaFetcheR was superior to existing approaches and databases by benchmarking the performance of the algorithm in two independent case studies based on two published datasets.

**Availability:** MetaFetcheR is available at https://github.com/komorowskilab/MetaFetcheR/.

## Introduction

Metabolomics allows the study of small molecule substrates and compounds that are involved in metabolic processes. Various complex diseases have been strongly linked to metabolic disorders, such as type 2 diabetes and cancer, making metabolomics a highly relevant field for single- and multi-omics studies (Diamanti *et al.*, 2019, 2020; Kaushik and DeBerardinis, 2018). Pathway enrichment analysis is a widespread analysis approach for metabolomics that require metabolites to map a predefined set of unique identifiers (Chong *et al.*, 2018). In this setup there are several issues that arise when accessing, pre-processing and analyzing metabolite data. For instance, the overlapping and non-overlapping information for metabolites is scattered across several small compound databases leading to major analysis and standardization issues (Wishart *et al.*, 2018; Hastings *et al.*, 2013; Kim *et al.*, 2019). Additional challenges occur with databases that deliver data with MySQL dump files or others that contain multiple entries for one metabolite or incomplete data. Finally, foreign reference identifiers may be missing, making it difficult, sometimes impossible, to find the link between two records of the same metabolite in different databases, while in other cases, the small fraction of reference identifiers that are present might lead to incorrect compounds. The aforementioned issues delay the delivery of results and more importantly, might lead to inconsistent or biased results.

Xia and colleagues have developed MetaboAnalyst, which is a versatile computational tool for metabolomics. This tool contains a module aimed at mapping names to identifiers of metabolites from the human metabolome database (HMDB), the chemical entities of biological interest (ChEBI), the Kyoto encyclopedia of genes and genomes (KEGG), PubChem and METLIN (Wishart *et al.*, 2018; Degtyarenko *et al.*, 2008; Hastings *et al.*, 2016; Kanehisa *et al.*, 2017; Kim *et al.*, 2019; Smith, 2005). However, the lack of a shared nomenclature for metabolite names commonly leads to numerous mismatches or no-matches. Additionally, Wishart and colleagues have mapped compounds of HMDB to identifiers of other databases that suffer from inconsistent matches (Wishart *et al.*, 2018). Moreover, the aforementioned tools map metabolite names to entries in HMDB that may lead to loss of information in case of a mismatch or absence of the metabolite from this specific database.

MetaFetcheR is a unified package targeted towards the metabolomics community that is able to resolve the multiple inconsistencies and incompleteness in data fetching. The algorithm exhaustively resolves such inconsistent cases and leads to improved mapping of small compounds to identifiers. This is showcased in two case studies using two published datasets (Diamanti *et al.*, 2019; Priolo *et al.*, 2014) and two existing mappers MS_targeted and MetaboAnalyst (Diamanti *et al.*, 2019; Chong *et al.*, 2018).

## MetaFetcheR algorithm

MetaFetcheR is an R package that uses the sparse input of primary database identifiers as a reference point to retrieve identifiers from other databases. The output of MetaFetcheR can be directly incorporated in analysis pipelines. The package unifies data from five open access and widely used small compound databases that include HMDB, ChEBI, PubChem, KEGG and Lipidomics gateway (LIPID MAPS) (Sud *et al.*, 2012). Each database has a standardized representation of the identifiers of compounds. The two most widely used representations that are also supported by MetaFetcheR, include the simplified molecular input line entry system (SMILES) (Weininger, 1988) and the IUPAC international chemical identifier (InChI) (Dashti *et al.*, 2017; Heller *et al.*, 2015) that describe chemical structures using ASCII characters.

The foundation of the underlying algorithm relies on constructing a local PostgreSQL database that acts as a cache memory of information. Initially, database dump files provided by HMDB, ChEBI and LIPID MAPS are to be downloaded. Subsequently, the bulk insertion function of MetaFetcheR is invoked to construct the local database (Supplementary Figure S1). In the interest of storage space and time we chose MetaFetcheR to cache data through HTTP calls on the fly from KEGG and PubChem. Cached instances from KEGG and PubChem are stored in the PostgreSQL database for later use in order to avoid unnecessary HTTP calls and timeouts due to excessive calls.

The input table to MetaFetcheR contains identifiers known by the user or provided by the metabolomics facility (Supplementary Table S1). The algorithm works on mapping the known identifiers of each metabolite to identifiers of other databases by filling in the empty fields. This is orchestrated via a queue-based algorithm. The algorithm handles exceptional cases of empty returns from a query by reiterating exhaustively until all identifiers have been retrieved or cannot be further resolved (Figure 1A). In addition, the algorithm allows storage of multiple mapped identifiers of the same compound from the same database that mark ambiguous situations. This allows the user to among the discovered identifiers choose the representative one that will be used in the downstream analysis. The ability of the algorithm to exhaustively resolve cases along with storing multiple discovered database identifiers for the same compound is one of the traits that allows MetaFetcheR to stand out. The algorithm optimizes speed performance by keeping track of formerly discovered records to avoid unnecessary iterations. A detailed presentation of the algorithm is presented in Supplementary Figure S2.

**Figure 1:**
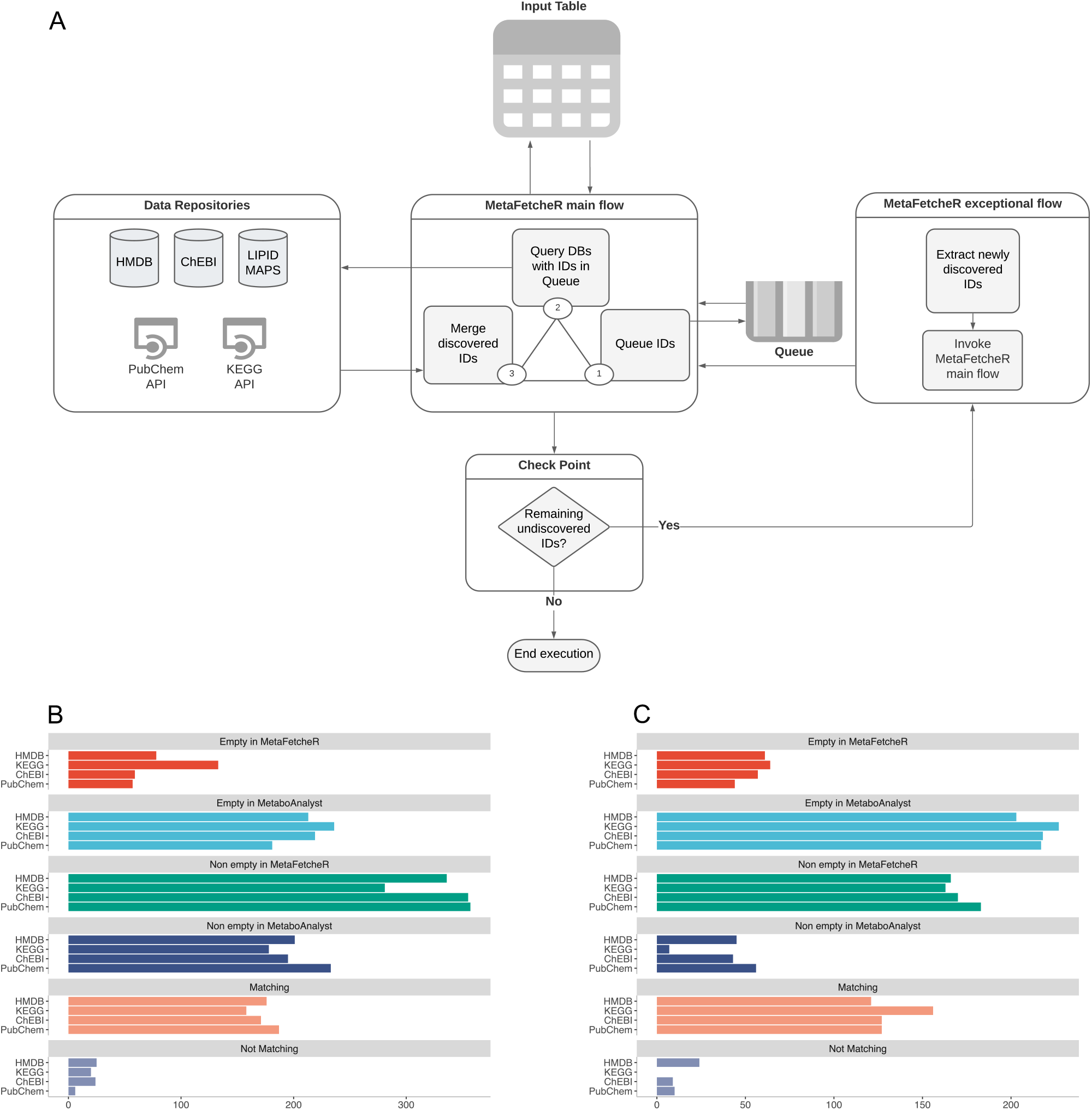
Overview of MetaFetcheR and comparison to other tools. **A)** A simplified graphic illustration of the MetaFetcheR algorithm. A detailed version is available in (Supplementary Figure S2). **B-C)** Comparison of the mapping performance of MetaFetcheR to MetaboAnalyst on the datasets from, **B)** (Diamanti *et al.*, 2019) and, **C)** (Priolo *et al.*, 2014). Empty in MetaFetcheR and MetaboAnalyst panels illustrate the number of identifiers that could not be mapped using the respective tool. Non-empty in MetaFetcheR and MetaboAnalyst panels present the number of identifiers that were successfully mapped using the respective tool. Matching panel shows the number of mapped identifiers that agreed between tools. Non-matching panel shows the number of mapped identifiers that were not in agreement between tools. The number of identifiers is shown on the x-axis.

### Usage scenarios and benchmarking

MetaFetcheR resolves problematic situations that arise when mapping identifiers of metabolites (Supplementary Figure S3). We benchmarked the matching performance of MetaFetcheR against two existing mappers in two case studies. We compared the performance of MetaFetcheR to MetaboAnalyst using two datasets from (Diamanti *et al.*, 2019; Priolo *et al.*, 2014). The mapping rate of MetaFetcheR was ~80% (non-empty fields), while MetaboAnalyst achieved ~48% mapping rate on (Diamanti *et al.*, 2019) (Figure 1B). For the dataset from (Priolo *et al.*, 2014) MetaFetcheR achieved ~78% non-empty fields rate, while MetaboAnalyst resulted in ~62% mapping rate (Figure 1C). A similar performance as for MetaboAnalyst was seen in the test utilizing MS_targeted (Supplementary Note - Benchmarking mapping performance of MetaFetcheR).

In addition to its competitiveness in mapping identifiers of metabolites, MetaFetcheR also provides insights into the quality of small compound databases. A test was run by selecting 1000 random identifiers from one of the five databases as input to MetaFetcheR and then we investigated the quality of the collection of retrieved identifiers. The test was performed 100 times for each database (Supplementary Tables S2-S6). The quality of the databases was assessed using three different metrics: *i)* percentage of consistency, *ii)* percentage of ambiguity, and *iii)* percentage of unresolved cases. KEGG showed the highest consistency percentage (~65%) and the lowest fraction of unresolved cases (~23%) compared to HMDB which had highest fraction of unresolved cases (~71%). Detailed results of the test and further explanation of the metrics can be found in Supplementary Note - Small compound databases quality test.

## Discussion

In this short report we presented MetaFetcheR, an R package for mapping metabolite identifiers across five small compound databases. Using two published datasets (Diamanti *et al.*, 2019; Priolo *et al.*, 2014) MetaFetcheR was shown to outperform other existing tools such as MS_targeted and MetaboAnalyst that provide similar mapping functionalities using. MetaFetcheR will be continuously updated to include additional databases. One of the important future updates includes the introduction of unique MetaFetcheR identifiers that will enhance the standardization of identifiers for small compounds.

## Author contribution

SY prepared the first draft of the manuscript, compiled scripts for the R package, designed benchmark strategies, generated results and assisted in the software and database design. RC wrote scripts to implement the algorithm and the database. KD is the owner of the main idea, co-wrote the manuscript and supervised the study together with JK.

## Acknowledgments

The authors thank Linda Holmfeldt and Aaron Vogan for reviewing the manuscript.

