## Supplementary notes and figures for "MetaFetcheR: An R package for complete mapping of small compound data"

#### Supplementary Material

### Supplementary Note

#### Algorithm detailed description

The input of the algorithm is in the form of a table with the relevant database identifiers for the five databases (HMDB, ChEBI, KEGG, LIPID MAPS, PubChem) (Wishart *et al.*, 2018; Hastings *et al.*, 2016; Kanehisa and Goto, 2000; Smith *et al.*, 2005; Kim *et al.*, 2019; Sud *et al.*, 2012). Database identifiers are stored in the columns while rows represent metabolites of interest whose identifiers require mapping (Supplementary Table S1). For each row the algorithm appends the known identifiers one by one to a queue. The algorithm fetches the db_id, where db is the database and id the identifier, from the top of the queue and queries the respective database for the record with this db_id as a primary identifier. The algorithm then fills all remaining identifiers for this metabolite with the returned results of the query. It might occur that the query return was empty; in which case the algorithm issues a new mapping attempt and queries the database again with the db_id as a secondary identifier, which several databases have. When the algorithm has filled in all the possible empty fields in each row that could be mapped using the primary or secondary identifiers that were initially given, it reiterates to check whether there are any remaining empty fields. In the case of an empty field, a reverse query is issued with one of the identifiers that were resolved during the first pass. The reverse query uses the discovered identifiers to query the respective database in an attempt to fill in the missing field. The algorithm reiterates until all identifiers have been filled in or cannot be further resolved. At the same time, it tracks already discovered records to avoid re-adding them to the queue. Linked identifiers are updated in the local database to avoid future queries and remapping. Moreover, the algorithm stores multiple mapped identifiers of the same compound from the same database that marks ambiguous situations. This allows the user to among the discovered identifiers choose the representative one that will be used for the downstream analysis.

The detailed flow chart of the discovery algorithm is shown in (Supplementary Figure S2). The input table provided by the user (df.input) is continuously updated during the execution of the algorithm. The output is in the form of an R data frame (df.result) that is merged with the input table (df.input) while the loop continues executing until the queue is empty. Upon exhaustion of the queue, reverse queries are executed. Once the algorithm has finalized, an updated R data frame similar to the input data frame structure is returned.

#### Small compound database quality test

The MetaFetcheR algorithm was able to provide insights into the quality of small compound databases as mentioned in the main text. The test was run by selecting 1000 random identifiers from one database (e.g., HMDB) to provide as input and investigate the quality of the collection of identifiers from other databases retrieved by MetaFetcheR. The test was performed 100 times for each database (Supplementary Tables S2-S6). We used three different metrics to assess the quality of each run: *i)* percentage of consistency, *ii)* percentage of ambiguity, and *iii)* percentage of unresolved cases. Consistency represents the percentage of one-to-one associated cases across all identifiers. Ambiguity is the percentage of original metabolite identifiers linked to multiple identifiers from other databases. Unresolved cases represent the percentage of cases that the original metabolite identifiers failed to link or were absent in all other databases. KEGG showed the highest consistency percentage (~65%), the largest fraction of ambiguous cases (~12%) and the lowest percentage of unresolved cases (~23%). HMDB showed the least ambiguity (~2%) and the highest number of unresolved cases (~71%) (Supplementary Figure S4).

#### Benchmarking mapping performance of MetaFetcheR

Performance of MetaFetcheR was benchmarked based on two case studies using two datasets and and two existing tools. The first dataset is by Diamanti et al. and the second one is by Priolo et al. The two tools are MS_targeted and MetaboAnalyst (Diamanti *et al.*, 2018; Priolo *et al.*, 2014; Chong *et al.*, 2018).

##### Case 1

We compared the performance of the algorithm for mapping metabolite identifiers to the mapped identifiers using MS_targeted along with the cases that were manually curated (that could not be mapped using MS_targeted). MS_targeted is a command line tool that was used for mapping metabolites identifiers. MS_targeted was run on Diamanti et al. dataset and had much higher number of unmapped identifiers (PubChem: ~99%, ChEBI: ~88%, LIPID MAPS: ~67%, KEGG: ~50% and HMDB: ~24%) compared to MetaFetcher (PubChem: ~14%, ChEBI: 14%, LIPID MAPS: ~67%, KEGG: ~32%, HMDB: ~19%). The results from MS_targeted were manually curated and compared to results of MetaFetcheR. There was ~80% match between MetaFetcheR mapped identifiers and MS_targeted mapped identifiers along with the manually curated identifiers that could not be mapped using MS_targeted (Supplementary Table S7 & Supplementary Figure S5). The results showed high concordance between the MetaFetcheR mapping and the manual curation of MS_targeted results which proves the superiority of MetaFetcheR over MS_targeted. The only database that had less than ~80% matches between mapped identifiers for both tools was LIPID-MAPS. This is possibly due to the sparse input for LIPID MAPS that had only 68 rows for LIPID MAPS. The used input dataset can be found in (Supplementary Table S8).

##### Case 2

We compared the MetaFetcheR mapping performance to that of the mapping function of MetaboAnalyst using data from (Diamanti *et al.*, 2018; Priolo *et al.*, 2014). For the comparison we only included metabolites that could be mapped by MetaboAnalyst. The MetaboAnalyst mapping function takes as input metabolite names that could be problematic since the name of the metabolites is not usually unique in all databases. For example, MetaboAnalyst could not map metabolites that had special characters at the end of their names. The list of metabolites that could not be mapped are listed in (Supplementary Table S9). Additionally, MetaboAnalyst could not map metabolites to the LIPID MAPS database identifiers. MetaFetcheR outperformed MetaboAnalyst on (Diamanti *et al.*, 2018) dataset (Figure 1B & Supplementary Table S10). MetaFetcheR had the lowest percentage of unmapped identifiers for all four databases (HMDB: 19%, KEGG: 32%, ChEBI: 14%, PubChem: 14%) compared to the unmapped identifiers using MetaboAnalyst (HMDB: 51%, KEGG: 57%, ChEBI: 53%, PubChem: 44%). Out of the metabolites that MetaboAnalyst managed to map (HMDB: 201 identifiers, KEGG: 178 identifiers, ChEBI: 195 identifiers, PubChem: 233 identifiers) MetaFetcheR did identify too (HMDB: 88%, KEGG: 89%, ChEBI: 88%, PubChem: 81%). MetaFetcheR had higher coverage of mapped identifiers (HMDB: 336 identifiers, KEGG: 281 identifiers, ChEBI: 355 identifiers, PubChem: 357 identifiers).

We compared the mapping performance of MetaFetcheR to MetaboAnalyst using (Priolo *et al.*, 2014) dataset. As mentioned previously, MetaboAnalyst takes metabolite names as input. Hence, we only included metabolites that could be mapped by MetaboAnalyst. There was only one metabolite that MetaboAnalyst could not map at all which was linolenate_[alpha_or_gamma_(18:3n3_or_6)]. MetaFetcheR outperforms MetaboAnalyst on

(Priolo *et al.*, 2014) dataset (Supplementary Table S11). MetaFetcheR had the lowest percentage of unmapped identifiers for all four databases (HMDB: 11%, KEGG: 0%, ChEBI: 4%, PubChem: 4%) compared to the unmapped identifiers using MetaboAnalyst (HMDB: 27%, KEGG: 28%, ChEBI: 25%, PubChem: 19%). Out of the metabolites that MetaboAnalyst could map (HMDB: 166 identifiers, KEGG: 163 identifiers, ChEBI: 170 identifiers, PubChem: 183 identifiers) MetaFetcheR could match with (HMDB: 73%, KEGG: 96%, ChEBI: 75%, PubChem: 69%). MetaFetcheR had higher coverage of mapped identifiers (HMDB: 203 identifiers, KEGG: 227 identifiers, ChEBI: 218 identifiers, PubChem: 217 identifiers) (Figure 2B & Supplementary Table S11). The dataset that was used as an input can be found in (Supplementary Table S12)

### Supplementary Figures

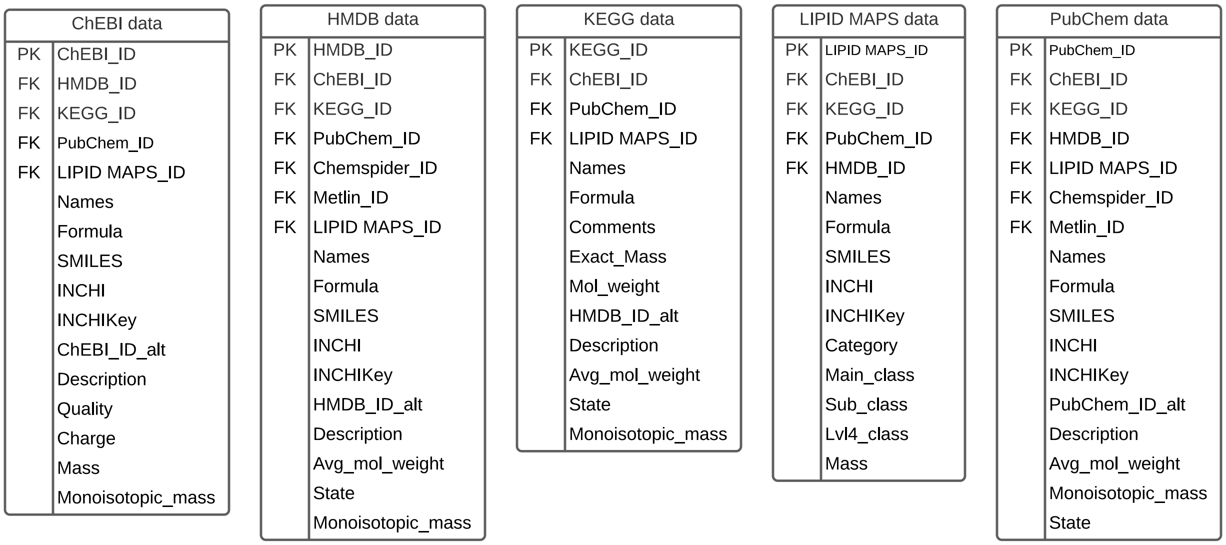


**Supplementary Figure S1:** Entity Relationship Diagram (ERD) of PostgreSQL database that MetaFetcheR builds locally. PK stands for primary key and FK stands for foreign keys.

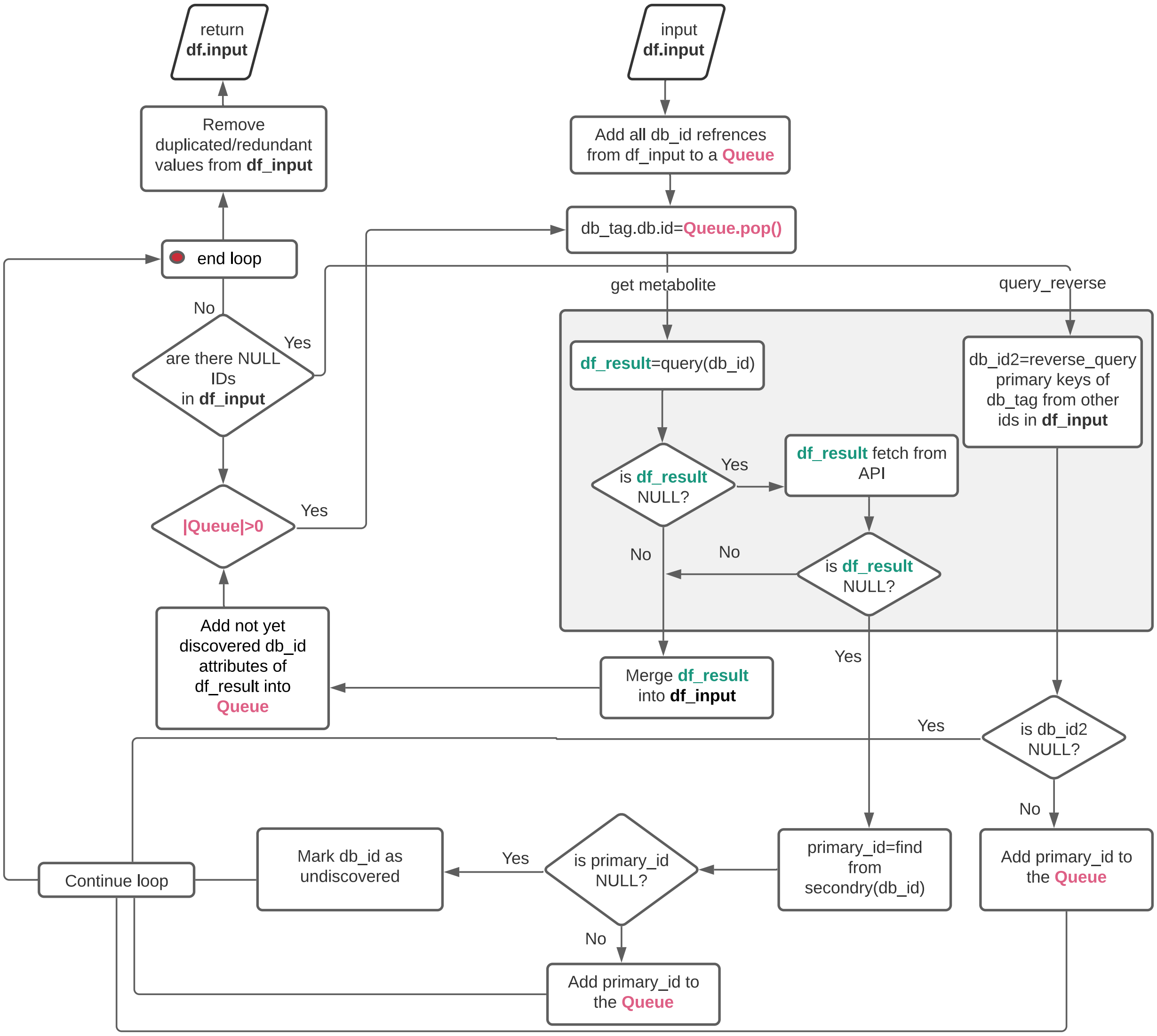


**Supplementary Figure S2:** Detailed flow chart of the MetaFetcheR algorithm. df.input is the initial table provided by the user and df.result is the output table. db_tag represents the name of the database, namely HMDB, ChEBI, KEGG, PubChem and LIPID MAPS, and db_id represents the id related to a certain db_tag. The queue is represented in bolded pink color, df_result, which represents the table that contains the query results, is represented in bolded green and df_input, which represents the input table, is represented in bolded black color. The grey box represents the main routine, which handles querying the different databases and APIs.

A

HMDB

| Metabolite Name | HMDB_ID | ChEBI_ID |
| --- | --- | --- |
| Diethyl_desulfide | HMDB0029572 | |

ChEBI

| Metabolite Name | ChEBI_ID | HMDB_ID |
| --- | --- | --- |
| - | - | |

B

HMDB

| Metabolite Name | HMDB_ID | ChEBI_ID |
| --- | --- | --- |
| ProstoglandinA1 | HMDB0002656 | 15545 |

ChEBI

| Metabolite Name | ChEBI_ID | HMDB_ID |
| --- | --- | --- |
| ProstoglandinA1 | 15545 | ? |

C

HMDB

| Metabolite Name | HMDB_ID | ChEBI_ID |
| --- | --- | --- |
| Fenoterol | HMDB0015405 | 149226 |

ChEBI

| Metabolite Name | ChEBI_ID | HMDB_ID |
| --- | --- | --- |
| Fenoterol | 149227 | HMDB0015405 |

**Supplementary Figure S3:** A representation of possible scenarios for mapping inconsistencies of metabolite identifiers between HMDB and ChEBI. The red color in the tables marks sample inconsistencies. **A)** The metabolite diethyl disulfide has an identifier in HMDB but does not have an entry in ChEBI. **B)** The metabolite prostaglandin A1 has an entry in HMDB with HMDB and ChEBI identifiers mapped to each other. However, the same metabolite has an entry in ChEBI but the link to HMDB is missing. **C)** The metabolite fenoterol has an entry in HMDB that is linked to ChEBI, however, the same metabolite has an entry in ChEBI with a different ChEBI identifier but the same HMDB identifier.

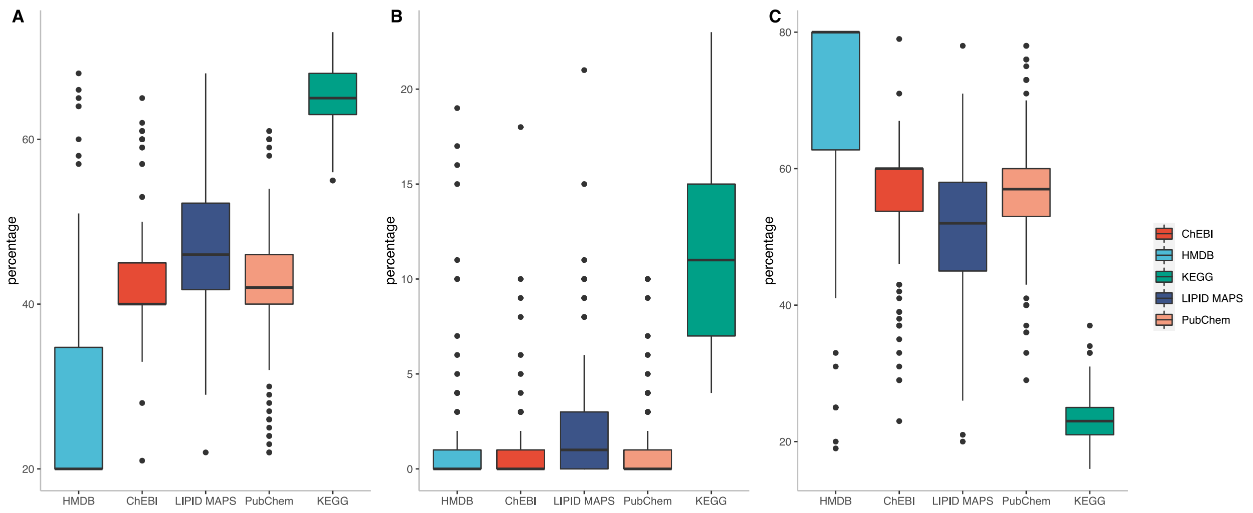


**Supplementary Figure S4:** The results for the test that was run on each database to investigate data quality. **A)** Box plot for the fraction of successfully mapped identifiers that had one-to-one mappings to the rest of database metabolite identifiers after running the test 100 times for each database. **B)** Box plot for the fraction of successfully mapped identifiers that had at least one one-to-many mappings to the rest of database metabolite identifiers after running the test 100 times for each database. **C)** Box plot for the extent of unmapped identifiers to the rest of database metabolite identifiers after running the test 100 times for each database.

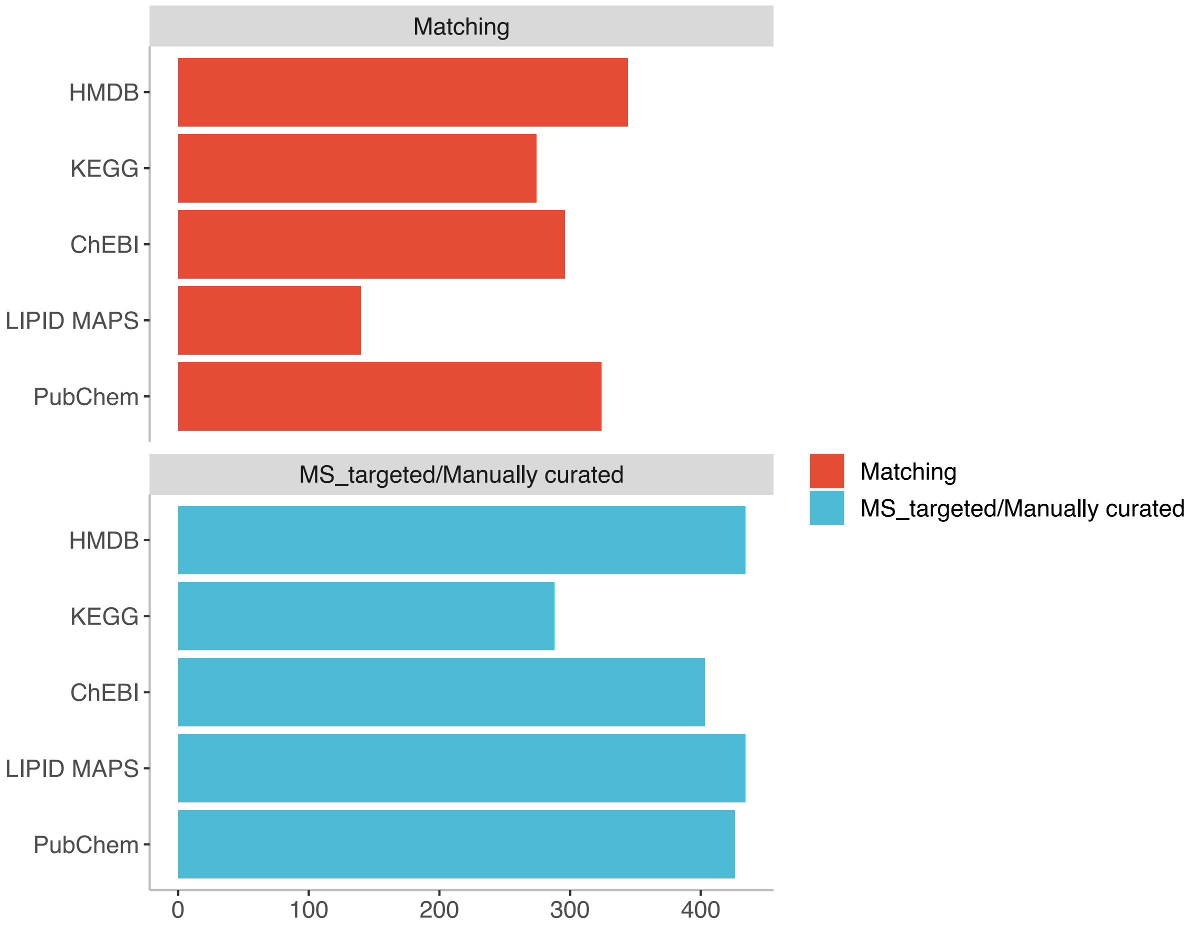


**Supplementary Figure S5:** Results of comparing MetaFetcheR to MS_targeted with manual curation. Red bars represent the number of identifiers mapped by MetaFetcheR that are in agreement with MS_targeted followed by manual curation. Blue bars represent the number of mapped identifiers by MS_targeted followed by manual curation that are in agreement with MetaFetcheR. There is ~80% overlap between MetaFetcheR mapped identifiers and MS_targeted mapped identifiers that have manually been curated.
